# Unveiling nesting strategies: Investigating visual concealment and nest-site selection in Yellow-wattled Lapwings, *Vanellus malabaricus* (Boddaert, 1783)

**DOI:** 10.1101/2023.08.13.552636

**Authors:** Vishwa Jagati, Geetha Ramaswami, Aniruddha Datta-Roy

## Abstract

Ground-nesting birds such as Yellow-wattled Lapwing, *Vanellus malabaricus* (Boddaert, 1783) employ several nesting and behavioral strategies to mitigate nest predation, which greatly influences their reproductive success and survival. Yet the breeding ecology of Yellow-wattled Lapwing has received relatively little research attention despite the species’ widespread presence in the Indian subcontinent. The objective of this study was to investigate the significance and impact of two visual concealment strategies commonly utilized by open ground-nesting birds: visual concealment through vegetative characteristics and camouflage using Yellow-wattled Lapwing as the model organism. We also assessed the nest-site selection of Yellow-wattled Lapwings in relation to vegetation cover and food availability, as well as their choice of nest substrate by using behavioral sampling, quadrat sampling, and digital image analysis techniques. We found that variation in hatching success could not be attributed to the analyzed camouflage mechanisms (disruptive markings and pattern complexity matching). Furthermore, the nesting pairs exhibited a preference for areas with high vegetation cover and low food availability while selecting nesting sites. We also showed that this species might exhibit an active behavioral preference for selecting nest substrates. It is important to note that this study was conducted during a single nesting season for the Yellow-wattled Lapwings, and further temporal replicates are necessary to validate and establish the findings over the long term.

## INTRODUCTION

Nest predation has been widely hypothesized as the primary factor contributing to the failure of reproductive success in ground-nesting birds (Ricklefs, 1969; Martin, 1993; Newton, 1998; Smith & Edwards, 2018). The selection of nest site (Burger et al., 1976), optimal clutch size (Martin, 1995), egg characteristics (Cott, 1940), and breeding behavior (Martin et al., 2000; Ghalambor & Martin, 2002) have been demonstrated to be influenced by predation pressure. Ground-nesting birds have developed several defensive adaptations against nest predation. A few such adaptations are behavioral defense, where they actively guard their nests and engage in aggressive behaviors to deter potential predators (Gochfeld, 1984; Kostoglou et al., 2020), nest concealment through vegetative characteristics (Götmark et al., 1995; Smart et al., 2006), and behaviorally induced camouflage (Summers & Hockey, 1981; Amat et al., 2012; Mayani-Parás et al., 2015; Troscianko et al., 2016). Additionally, some species have evolved to use cryptic nest sites (Lovell et al., 2013; Gómez et al., 2018) or exhibit cryptic egg characteristics, such as egg coloration or pattern that matches the surrounding environment (Wallace, 1889; Cott, 1940; DeLong et al., 1995; Skrade & Dinsmore, 2013; Stoddard et al., 2016).

Nesting birds are expected to choose nest sites that maximize their reproductive success (Gjerdrum et al., 2005). In ground-nesting species, nest-site selection is commonly influenced by factors such as the risk of nest predation, nest substrate, vegetative characteristics, and availability of foraging substrates (Gómez et al., 2018; Laidlaw et al., 2020; Momberg et al., 2023). Birds often choose nest sites that minimize the risk of predation, as this can greatly affect their hatching success (Alonso et al., 1991; Tuomenpuro, 1991; Collias & Collias, 2014). Topography plays a crucial role in the nest-site selection for many bird species, particularly ground-nesting, as it provides the ability to either detect potential predators from a distance or remain concealed in the vegetation’s cover to avoid detection (Muir & Colwell, 2010; Colwell et al., 2011; Cunningham et al., 2016; Korne et al., 2020; Momberg et al., 2023). Visual concealment of the nests through egg characteristics (Egg camouflage) and vegetation cover are frequently observed among ground-nesting birds and are likely to affect their vulnerability to predation (Laidlaw et al., 2020). Certain bird species, particularly waders, employ a nesting strategy where nests are constructed on bare ground, and the adult plumage and/or egg coloration blend in with the surrounding environment, providing camouflage (Troscianko et al., 2016; Laidlaw et al., 2020). Breeding in open landscapes allows these bird species to rely on early predator detection (Amat & Masero, 2004), and lower vegetation cover can result in an early departure from nests upon detecting potential predators (Gómez-Serrano & Lopez-Lopez, 2014). On the other hand, waders may also prefer nest sites that allow for the concealment of both nests and incubating adults within the surrounding vegetation (Götmark et al., 1995). This strategy may involve selecting sites with dense vegetation. However, incubating parents hidden in dense vegetation can be at a disadvantage as they may not be able to detect predators early through visual cues (Laidlaw et al., 2020).

Additionally, the availability of food resources, which is intricately linked to vegetation, is a vital factor that can substantially influence nest site selection by birds (Li & Martin, 1991; Momberg et al., 2023). Parental effort during incubation is another crucial factor that can determine reproductive success in birds (Matysiokova & Remeš, 2010). During the incubation period, it is common for parents to forage at considerable distances away from their nest, resulting in the nest being vulnerable to potential predation (Eikenaar et al., 2003; Vafidis et al., 2018). Therefore, parental birds must make a trade-off between the time they spend foraging away from the nest and the time they spend at the nest performing tasks such as incubation, guarding, and defending against potential predators (Orians, 1979; Martin, 1992). However, if there is ample availability of food near the nest, it is expected to shift the trade-off in favor of increased time spent on the nest for incubation and prompter defense against potential predators (Vafidis et al., 2018). Hence, comprehending how food availability, vegetation cover, and predation pressure are interconnected with nest-site selection is essential in the study of bird behavior and ecology.

The evolutionary influence of nest predation on the appearance and pigmentation of bird eggs, particularly in the order Charadriiformes, serves as a defense mechanism against visually-oriented predators, highlighting its vital role in shaping avian reproductive strategies (Cott, 1940; Hockey, 1982; Solis & De Lope, 1995; Lloyd et al., 2000; Kilner, 2006). It has been postulated that the adaptive value of egg characteristics lies in their ability to provide effective camouflage, aiding in the survival of bird eggs (Wallace, 1889; Skrade & Dinsmore, 2013). The influence of bird egg appearance on nest success has been a persistent area of research interest for evolutionary biologists, with notable scientists like Alfred Wallace (Wallace, 1889), Hugh B. Cott (Cott, 1940), and David Lack (Lack, 1958) contributing to our understanding of this phenomenon. Recent years have witnessed significant progress among scientists in understanding the diverse forms of camouflage, evaluating their survival advantages, and unraveling the underlying mechanisms related to visual processing (Stevens, 2013). Yet, limited research has been conducted to investigate the influence of nest and egg characteristics on clutch survival (Hockey, 1982; Solis & De Lope, 1995; Westmoreland & Kiltie, 1996; Lloyd et al., 2000; Nguyen et al., 2007; Lee et al., 2010; Colwell et al., 2011; Lovell et al., 2013; Troscianko et al., 2016; Stoddard et al., 2016), and most studies in this field have primarily focused on the colors of eggs and background substrate, rather than more intricate aspects such as egg surface patterns and background substrate textures (Underwood & Sealy, 2002; Stoddard et al., 2016).

The amount and arrangement of maculation or markings on eggshells, as well as the textures of the background substrate, are crucial factors that can affect egg camouflage and potentially affect nest survival (Stoddard et al., 2016). In spite of that, only a handful of studies have explored the characteristics of egg markings or background substrate textures in relation to nest survival (Westmoreland & Kiltie, 1996; Nguyen et al., 2007; Troscianko et al., 2016; Stoddard et al., 2016). Background matching and disruptive coloration are among a variety of concealment techniques that have been proposed for effective camouflage (Thayer, 1918; Cott, 1940; Stevens & Merilaita, 2009a). In disruptive coloration, highly contrasting outline patterns of an organism break up its shape against the background and conceal the organism’s outline (Kang et al., 2015; Stoddard et al., 2016). It functions by inhibiting the detection of an organism’s true shape and outline (Stevens & Merilaita, 2009b; Kang et al., 2015). In background matching, an organism achieves camouflage by possessing body colors and/or patterns that match those in its background substrate (Stoddard et al., 2016).

To blend in with their surroundings and remain undetected, camouflage often involves matching the visual appearance of the background. One effective way to optimize camouflage is to select optimal backgrounds that complement an individual’s appearance (Gómez et al., 2018). If there is variation in nest substrate preference among individuals of a single species, the behavioral selection of appropriate substrates and backgrounds could be a significant strategy for enhancing camouflage and improving nest success (Stevens et al., 2017). It is currently uncertain if females possess the genetic ability to modify the coloration and patterns of their eggs to match their surroundings (Gosler et al., 2000). However, an alternative explanation proposed in the study (Lovell et al., 2013) suggests that females may select microhabitats based on the appearance of their eggshells, as demonstrated in laboratory settings with Japanese quail. To confirm this hypothesis, further research conducted in natural environments is needed, taking into account multiple selective factors that can influence camouflage (Gómez et al., 2018).

Due to their unique ecological and ground-dwelling characteristics, ground-nesting birds are an ideal model system for conducting behavioral studies. Yellow-wattled Lapwings present an ideal species for conducting such studies. Interestingly, two distinct types of nests have been observed in this species: nests constructed using pebbles, stones, or soil balls, and nests constructed using grass materials or with eggs laid in the middle of grassy patches (Greeshma & Jayson, 2018) (Figure 1). One particularly compelling aspect is the study of camouflage mechanisms in ground-nesting bird eggs in natural settings, for two main reasons. Firstly, clutch predation can significantly impact their reproductive success (Macdonald & Bolton, 2008), making camouflage through egg coloration and patterning an important defense mechanism against predator detection. Secondly, photographing egg clutches in their natural habitats and documenting their survival is relatively easy, enabling the assessment of egg camouflage in wild populations without artificial stimuli or lab settings (Stoddard et al., 2016).

**Figure 1.**
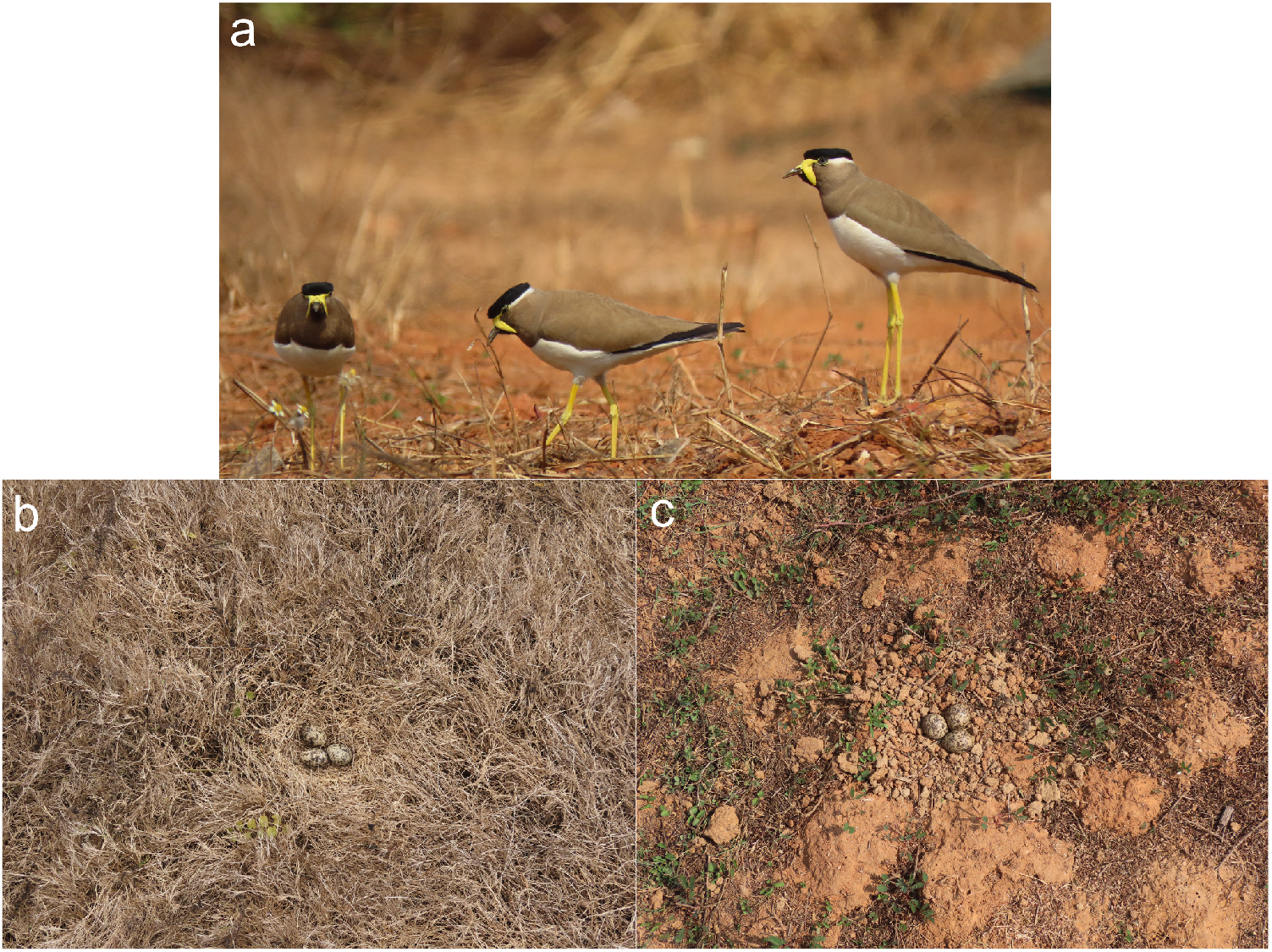
(a) Yellow-wattled Lapwings in their natural habitat, and types of Yellow-wattled Lapwing nests: (b) grass nest, (c) stone nest.

Our study had four primary goals. First, to analyze the nest-site selection in Yellow-wattled Lapwings focusing on the factors of food availability and vegetation cover. Second, to quantify the two major visual concealment strategies adopted by these ground-nesting birds against nest predation, (i) nest concealment through vegetative characteristics, and (ii) egg camouflage. Specifically, we investigated pattern complexity matching (a form of background matching) and disruptive effects (a form of disruptive coloration). Our third goal involved examining the relationship between the hatching success of clutches and the availability of food at nest sites, alongside the nest concealment through vegetation cover and egg camouflage. Lastly, we sought to investigate whether Yellow-wattled Lapwings actively select nest substrates to camouflage their eggs. We conducted a comparative analysis of different nest types, distinguishing between those constructed with pebbles, stones, or soil balls and those made with grass materials or with eggs laid in the middle of grassy patches, to determine the camouflage mechanisms utilized.

## METHODS

### STUDY SPECIES

The Yellow-wattled Lapwing is an elegantly long-winged plover with brown plumage, and distinct yellow wattles and legs (Figure 1) (Grimmett et al., 2016). It is endemic to the Indian subcontinent and can be commonly observed across the entire region (Ali & Ripley, 1983; Mukherjee et al., 2015; Grimmett et al., 2016). Yellow-wattled Lapwings breed during the dry season, typically between March and May in India (Jayakar & Spurway, 1965a). The coloration and markings of Yellow-wattled Lapwing eggs are well-adapted for camouflage (Gupta et al., 2022). The eggs are typically pale brown to sandy-colored, with darker speckles or spots (Ali & Ripley, 1983). Their nest is a simple scrape on the ground, often lined with pebbles, vegetation, or other materials that blend with the surroundings (Ali & Ripley, 1983). There are two observed types of nests - those constructed using pebbles, stones, or soil balls, and those made of grass materials or with eggs laid in the middle of grassy patches (Greeshma & Jayson, 2018) (Figure 1). The clutch size of Yellow-wattled Lapwings is usually 3-4 eggs (Jayakar & Spurway, 1965a; Ali & Ripley, 1983). They primarily feed on spiders and insects such as beetles, termites, grasshoppers, and crickets (Jerdon, 1864). Despite being a fascinating species, research on Yellow-wattled Lapwings is limited and mostly comprises natural history notes (Jayakar & Spurway, 1965a, 1965b, 1968; Dhindsa, 1983; Sethi et al., 2010; Chand Gupta & Kumar Kaushik, 2012; A. Kumar & Kanaujia, 2015; C. Kumar, 2015; Mukherjee et al., 2015; Chavan et al., 2016; Greeshma & Jayson, 2018).

### STUDY SITE

The study was carried out within the premises of the National Institute of Science Education and Research (NISER) campus, Bhubaneswar, Odisha, India (Figure 2). NISER is located close to the eastern coast of India, situated within a region characterized by a tropical savanna climate (Moharana et al., 2022). The campus vegetation predominantly consists of seasonal grasslands and plantations. The collection of field data occurred from December 2020 to February 2022. Two distinct locations within the campus were selected for the study: the football ground and the cricket ground, which offered two different habitats for ground-nesting birds (Figure 2).

**Figure 2.**
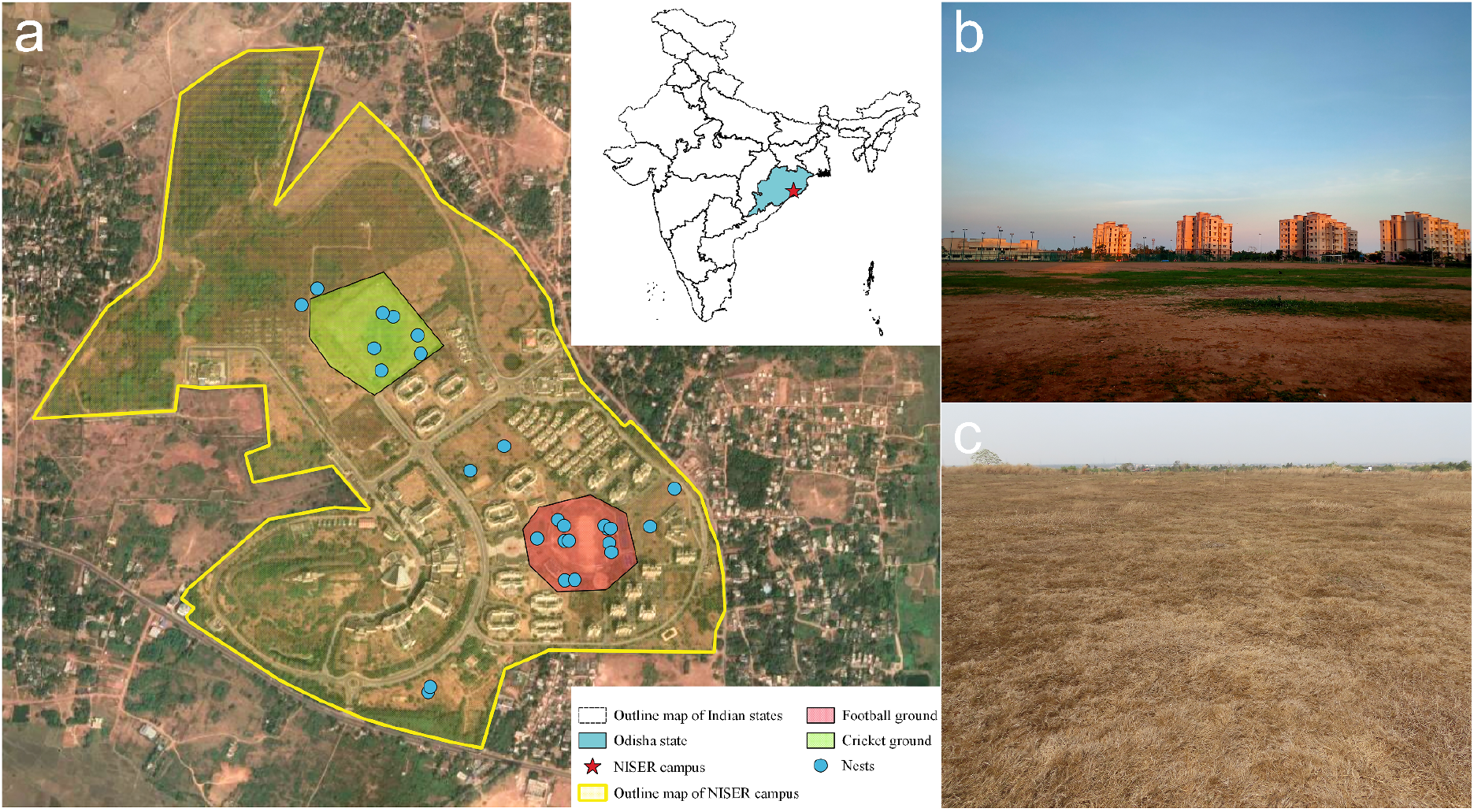
(a) Map depicting the study site, targeted habitats, and the location points of observed nests; (b) the football ground, and (c) the cricket ground.

The football ground is characterized by moderate vegetation cover and is subject to frequent human disturbance due to recreational activities. Additionally, it has a relatively high population of feral dogs (N∼15). High-intensity artificial light sources are used to illuminate this ground daily. In contrast, the cricket ground is an abandoned open area with high vegetation cover, very little human disturbance, and a smaller population of feral dogs (N∼5). No floodlights are present in this location. The study area’s common nest predators include feral dogs, feral cats, Indian jungle crows (*Corvus culminatus*), shikras (*Accipiter badius*), black kites (*Milvus migrans*), black-winged kites (*Elanus caeruleus*), and barn owls (*Tyto alba)*.

### NEST-SITE CHOICE AND HATCHING SUCCESS

We used direct observations of individuals to understand adult birds’ nest-site choice and the hatching success of young birds. The study involved monitoring Yellow-wattled Lapwings for a fixed period of six hours each day (from 6:30 am to 9:30 am and 3 pm to 6 pm) at two study sites on alternate days from December 2020 to February 2022. A total of approximately 1800 man-hours were spent on field observations. Observations were made from a distance using a pair of Nikon Aculon a211 10×50 binoculars. Nests were located based on the behavior of the parent birds, such as scraping the ground for nesting, incubating, or threatening predators. A total of 26 nests were found during the study period. Among these, 12 nests were found on the football ground, while 8 nests were observed on the cricket ground. The remaining nests were located outside of these two habitats. Nests were monitored every 2-3 days until 25 days into incubation or until pipping and/or noises from the chicks were observed, after which they were checked daily until hatching success data was obtained. Despite being laid within a time interval of a few days, the eggs exhibit synchronous hatching. Hatching success for a nest was determined by the successful hatching of at least one egg, even if the incubation process was not optimal for the other eggs. Hatching failure could occur due to parental abandonment, anthropogenic activities, ineffective incubation, or predation.

We used images of nests with complete clutches to quantify egg camouflage. The images of the nests were captured from about one meter above the ground using a Canon SX70hs. The images of completed clutches (taken five days after incubation initiation) were selected for image analysis (Stoddard et al., 2016), and all images were saved in RAW (.CR3) file format.

### VEGETATION COVER AND FOOD AVAILABILITY

We used quadrat sampling to assess the availability of food and vegetation cover at nest sites (Gleason, 1920; Gardiner et al., 2005) to understand how these factors affected the selection of nest sites by adult birds. This involved placing a quadrat with an area of 10 m^2^ around the nest, and within this larger area, five quadrats with an area of 1 m^2^ were placed randomly. The decision to use a 10 m^2^ quadrat for analyzing nest-site choice was based on the observation that parents tended to exhibit defensive behavior against the potential predators within this approximate range. One possible explanation for this behavior is that the visual concealment provided by vegetation might become less effective within this range. It was also noted in most instances that at least one parent remains within this area while foraging. Each 1m^2^ quadrat within the 10m2 plot was subdivided into 25 smaller plots of area 20cm^2^. The number of insects and the smaller plots intersecting with vegetation were then counted in these 1m^2^ quadrats, and this data was used to estimate vegetation and insect density.

### IMAGE ANALYSIS

We utilized digital image analysis techniques and edge detection algorithms to quantitatively investigate how Yellow-wattled Lapwings employ camouflage to protect their eggs. To maintain consistency in the analysis, photographs taken at the same time of day were chosen. Additionally, to standardize the images, the converted files from RAW to TIFF format were processed using PictoColor inCamera software (iCorrect EditLab Pro), which included equalization and linearization adjustments (Stevens et al., 2007). This procedure ensured that the images underwent a standardized transformation, allowing for accurate and comparable analysis across the dataset. We followed methods described by Stoddard et al., (2016) to quantify aspects of camouflage. We used similar variable names where possible. Our first step was to divide the image into three sections: the egg contour region (the outer edges of the clutch), the internal egg region (the surface area of the clutch of eggs), and the background substrate region (the ground substrate surrounding the clutch) (Figure 3). We then quantified variables related to the focused forms of camouflage: pattern complexity matching, and disruptive effects. In terms of pattern complexity matching, animals exhibit enhanced camouflage when their resting or background substrate possesses more complex patterns than their own body patterns. This allows them to blend in effectively with their surroundings by utilizing the complexity of the environment. In disruptive effects, animals employ strategies to conceal their true outline by leveraging the patterns present in the background substrate or their own body patterns. By manipulating these elements, they can disrupt the perception of their distinct shape, making it harder for predators to detect them. Our analyses were coded in MATLAB (MathWorks Inc. 2022), following Stoddard et al. (2016).

**Figure 3.**
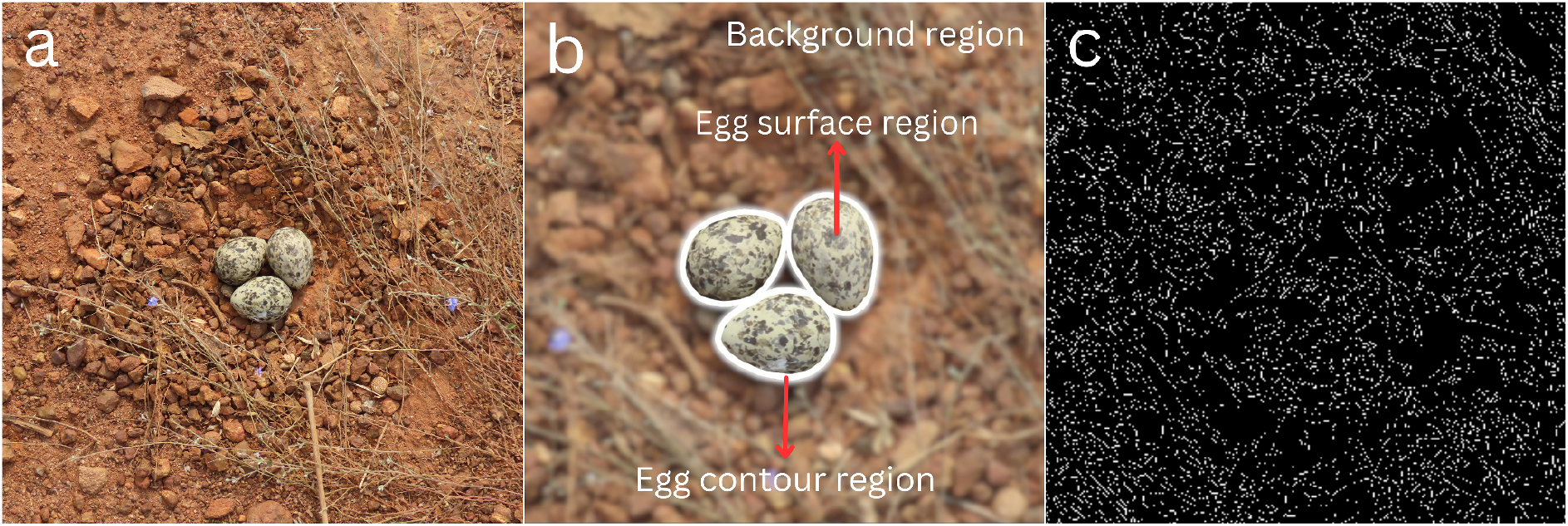
(a) A Yellow-wattled Lapwing nest; (b) Each nest image was separated into 3 sections for image analysis-background substrate region, internal egg region, and egg contour region (zoomed in); (c) An illustrative image demonstrating the application of edge detection algorithm to detect the edges in both the eggs and the background substrate based on Figure 3a and 3b (zoomed-in).

The Canny edge detection algorithm was used to identify edges in the images using the MATLAB Image Processing Toolbox. The algorithm was configured with a threshold of 0.2 and a sigma of 3 (Stoddard et al., 2016). This computer vision algorithm is widely used and works by searching for local maxima of the intensity gradient in the image to locate edges (Canny, 1986). While most research on the detection of edges has focused on humans, other vertebrate visual systems may employ similar mechanisms to detect the edges (Stoddard et al., 2016).

### CAMOUFLAGE QUANTIFICATION

#### Quantifying pattern complexity matching

Animals can enhance their camouflage by selecting resting surfaces with intricate patterns, where the background substrate exhibits more complex patterns than the animal’s markings. This strategy can effectively improve camouflage irrespective of the animal’s own pattern (Merilaita, 2003). Additionally, an animal can further enhance its camouflage by having a pattern that matches the background pattern (Dimitrova & Merilaita, 2014). To quantify the level of complexity in the background pattern, we measured the proportion of edges detected in the background substrate region (background edges). To measure the match between an egg surface maculation and the background, we calculated the complexity ratio (Table 1). In theory, a lower complexity ratio would result in better camouflage because it would indicate that the edges of the background are more prominent compared to the edges of the egg surface (Merilaita, 2003). Additionally, when the complexity ratio is closer to 1, it suggests that the organism has a higher degree of pattern complexity matching (Merilaita, 2003; Stoddard et al., 2016).

**Table 1.** Variables of camouflage mechanisms: Pattern complexity matching and Disruptive effects.

| Camouflage mechanism | Variables | Definition |
| --- | --- | --- |
| Pattern complexity matching | Background edges (BgEdge) | The ratio of edge pixels in the background substrate and the total number of pixels in the background substrate |
|  | Egg edges or Internal edges (EggEdge) | The ratio of edge pixels in the egg surface region and the total number of pixels in the egg surface region |
|  | Complexity ratio (CompRat) | The ratio of EggEdge and BgEdge |
| Disruptive effects | Contour edges (ContEdge) | The ratio of edge pixels in the egg's contour region and the total number of pixels in the egg contour region |
|  | Disruptive ratio (DisRat) | The ratio of ContEdge and EggEdge |
|  | Visibility ratio (VisRat) | The ratio of ContEdge and BgEdge |

### Quantifying disruptive markings

Organisms can also achieve camouflage by using disruptive markings to hide their edges. It occurs when the egg surface patterns or substrate patterns make it difficult to detect the organism’s outline. To quantify the degree of edge hiding, we calculated the proportion of edges in the egg contour region (Contour edges), the disruptive ratio, and the visibility ratio (Table 1) (Stoddard et al., 2016). When the disruptive ratio and visibility ratio are less than 1, the edges in the contour are harder to detect than those in the eggs/background substrate, resulting in low detectability of the eggs. Conversely, when they are greater than 1, the edges in the contour are easier to detect than those in the eggs/background substrate, leading to high detectability of the eggs (Stoddard et al., 2016). Therefore, the lower the disruptive and visibility ratios, the better would be the camouflage.

### STATISTICAL ANALYSES

The objective of our study was to investigate the influence of vegetation cover and food availability on the nest-site selection of Yellow-wattled Lapwings. Additionally, we explored how camouflage variables impacted the survival of the clutches. A comparative analysis of camouflage between two observed nest substrate choices, grass, and stone, was also conducted, focusing on pattern complexity matching and disruptive effects.

For investigating the relationship between the number of nests found and factors such as vegetation cover and food availability, we utilized a Generalized Linear Modeling framework with a Poisson error structure for vegetation cover and a Quasi-Poisson error structure for insect density. Furthermore, we examined the impact of key factors, including vegetation cover, food availability, and the variables of pattern complexity and disruptive effects, on the hatching success of the lapwings’ eggs using a Generalized Linear Model (GLM) framework with a binomial error structure. We assessed parameter estimate significance through bootstrapping – with the assumption that estimates with 95% CI not overlapping zero, have a significant effect on the response variable.

We assessed the normality of errors using the Shapiro-Wilk test and homogeneity of variance using the F-test of the different camouflage metrics. Based on these assessments, we compared camouflage metrics across different nest substrates using a t-test for normally distributed errors (for BgEdge, EggEdge, ContEdge, DisRat, and VisRat), and a Mann Whitney U test for non-normally distributed errors (for CompRat).

All statistical analyses were run in R version 3.6.1 (R Core Team 2019).

## RESULTS

### NEST-SITE CHOICE AND HATCHING SUCCESS

Yellow-wattled Lapwings in the study area nested between November and June/July – a seasonal pattern that seems to be earlier in the year than that reported from other studies (Ali & Ripley, 1983; Sethi et al., 2010; C. Kumar, 2015). Based on our analyses, vegetation cover seems to predict the abundance of nests reasonably well (Pearson’s R of observed vs model predicted values = 0.83) (Table 2, Figure 4a). However, bootstrapped 95% CI for this parameter overlapped 0 suggesting that the coefficient of vegetation cover may not be statistically significant, and the effects of this variable may be more apparent with increased sampling effort. On the other hand, insect density was a statistically significant predictor of the number of nests (Table 2, Figure 4b).

**Table 2.** Regression results for hatching success and number of nests in relation to various variables.

| Predictor variable | Response variable | Bootstrap value |  | Estimated coefficient | z-value | Pearson's R |
| --- | --- | --- | --- | --- | --- | --- |
|  |  | Lower 95% CI | Upper 95% CI |  |  |  |
| Vegetation cover | Hatching success | -4.335 | 1.992 | 0.005 | 0.335 | NA |
| Insect density | Hatching success | -0.572 | 2.654 | -0.297 | -1.434 | NA |
| Complexity ratio | Hatching success | -7.316 | 1.296 | 3.303 | 1.636 | NA |
| Disruptive ratio | Hatching success | -3.792 | 10.26 | -2.800 | -1.158 | NA |
| Visibility ratio | Hatching success | -8.742 | 3.854 | 2.247 | 1.16 | NA |
| Vegetation cover | Number of nests | -1.113 | 0.430 | 0.022 | NA | 0.83 |
| Insect density | Number of nests | 1.720 | 2.638 | -0.258 | NA | 0.9 |

**Figure 4.**
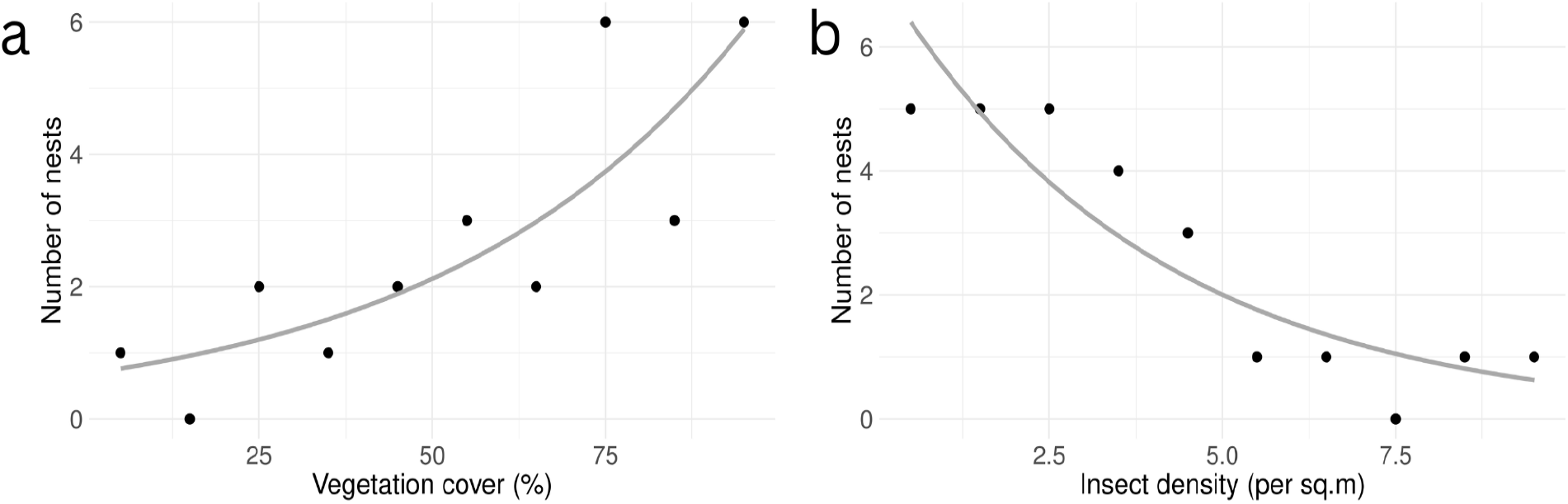
Correlation of (a) vegetation cover, and (b) insect density with number of nests.

The relationship between hatching success and vegetation cover was inconclusive, with a non-significant estimate (Table 2, Figure 5a). Interestingly, we observed a trend that nests with lower food availability in their surroundings had relatively higher hatching success rates (Figure 5b). However, this relationship was not statistically significant as the 95% confidence interval for the insect density coefficient includes zero (Table 2), which could be due to the low sample size. Based on the available data, we lack evidence to conclude that food availability or vegetation cover significantly affects hatching success.

**Figure 5.**
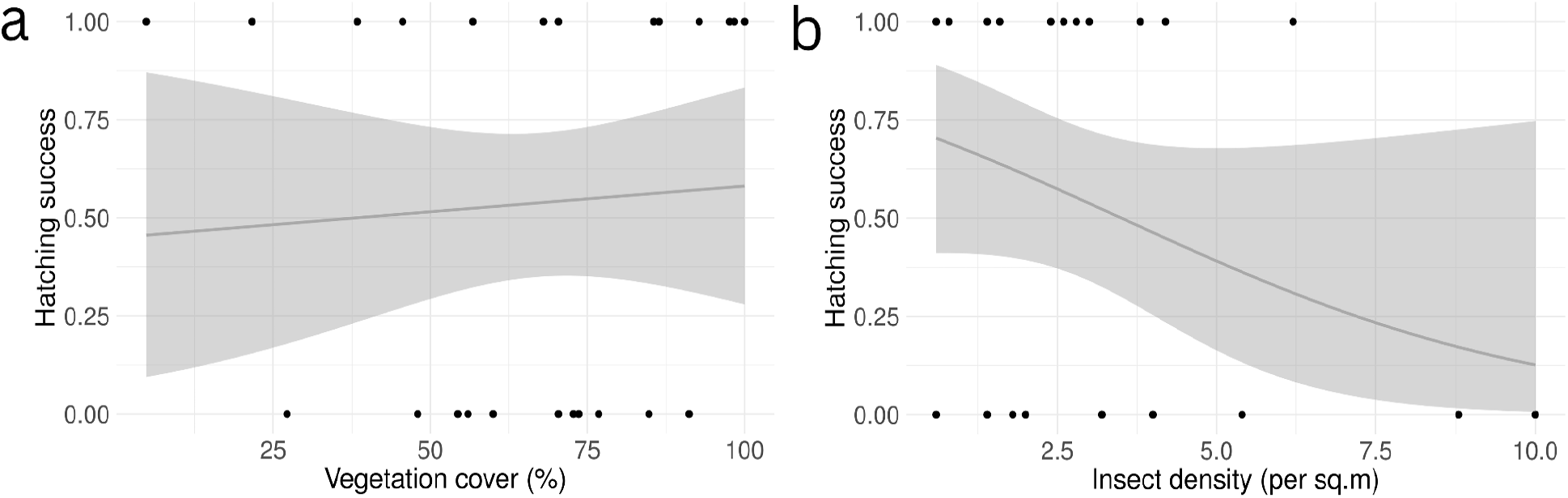
Hatching success in relation to (a) vegetation cover and (b) insect density (shaded region along the lines represents the 95% confidence level).

### CAMOUFLAGE AND HATCHING SUCCESS

We found no evidence of significant associations between hatching success and complexity ratio, disruptive ratio, or visibility ratio (Table 2). However, we report here some of the broader trends observed in the data which may be ecologically relevant: higher complexity ratios were positively associated with successful hatching, aligning with theoretical expectations (Figure 6). This finding suggests that nests laid on background substrates with relatively fewer edges than egg surface edges experienced enhanced hatching success. Contrary to theoretical expectations, where background substrates with more edges than eggs’ surface edges are believed to achieve better camouflage, our study demonstrated a different outcome. Consistent with theoretical predictions, our investigation indicated that nests with lower disruptive ratios exhibited higher hatching success rates. As the disruptive ratio increased, the likelihood of successful hatching decreased (Figure 7a). In contrast, our study’s findings regarding the visibility ratio contradicted theoretical expectations. The theory proposes that nests with lower visibility ratios should possess better camouflage and experience higher hatching success rates. However, our data showed the opposite result: an increase in visibility ratio corresponded to an increase in the probability of successful hatching (Figure 7b).

**Figure 6.**
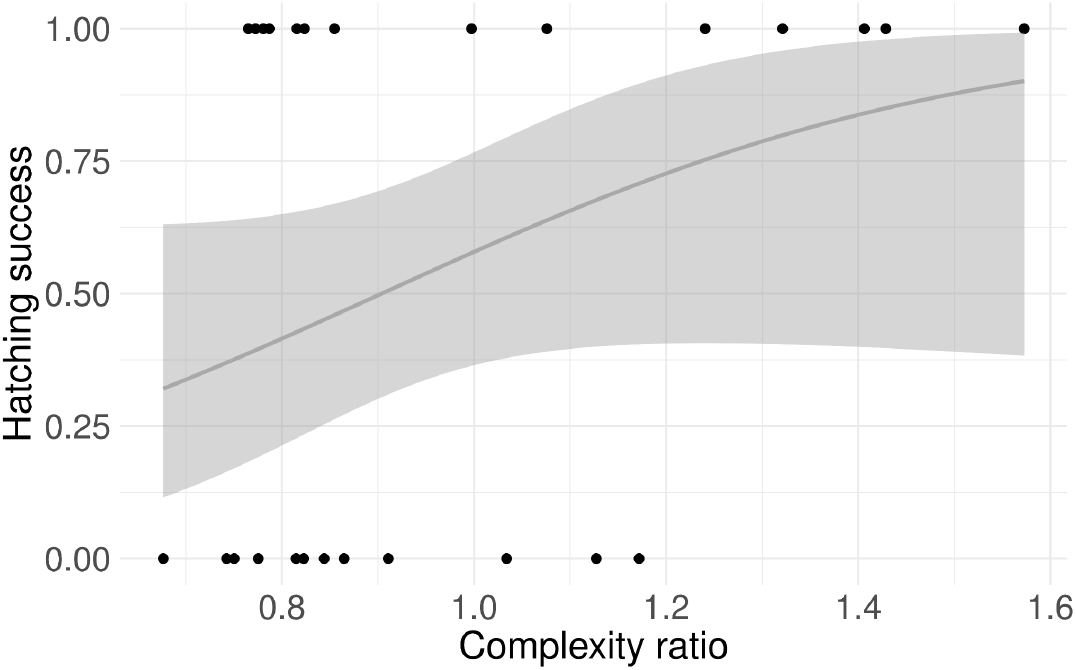
Hatching success in relation to the complexity ratio (shaded region along the lines represents the 95% confidence level).

**Figure 7.**
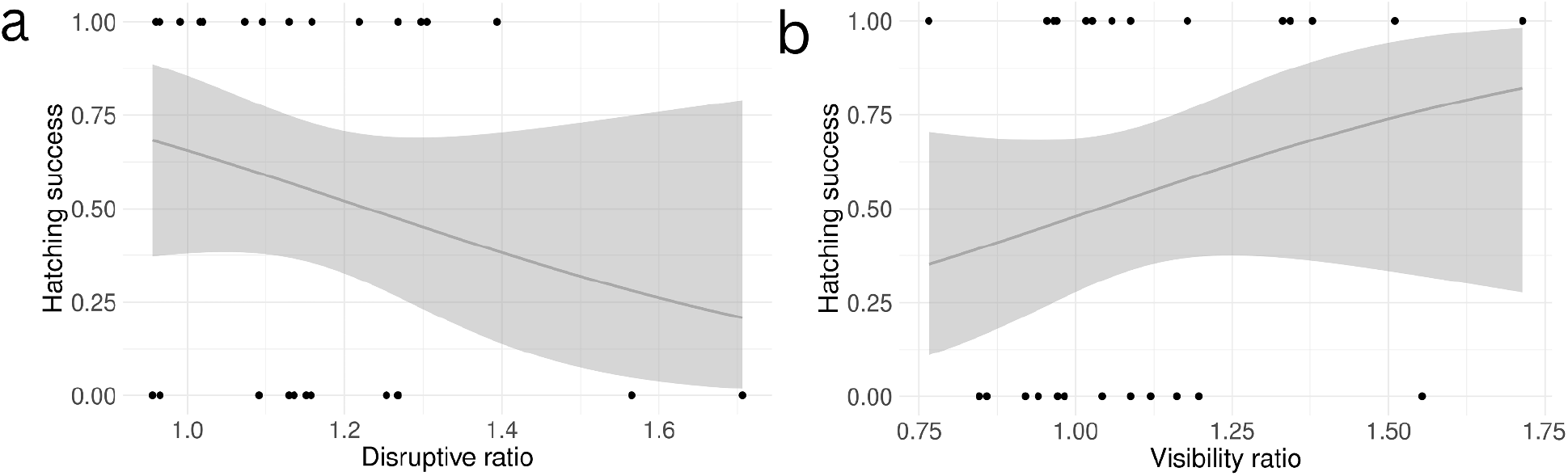
Hatching success in relation to (a) disruptive ratio and (b) visibility ratio (shaded region along the lines represents the 95% confidence level).

### NEST SUBSTRATE CHOICE - GRASS NESTS VS STONE NESTS

#### Comparing pattern complexity matching

Grass nests were constructed on substrates with higher edge complexity, indicating potentially better camouflage compared to stone nests based on this metric (Merilaita, 2003; Stoddard et al., 2016) (Figure 8). In terms of the number of surface egg edges (EggEdge), no significant difference was observed between the two nest types (p-value=0.228; t-value=1.246). This suggests that both grass and stone nests exhibit a similar level of surface egg edge complexity.

**Figure 8.**
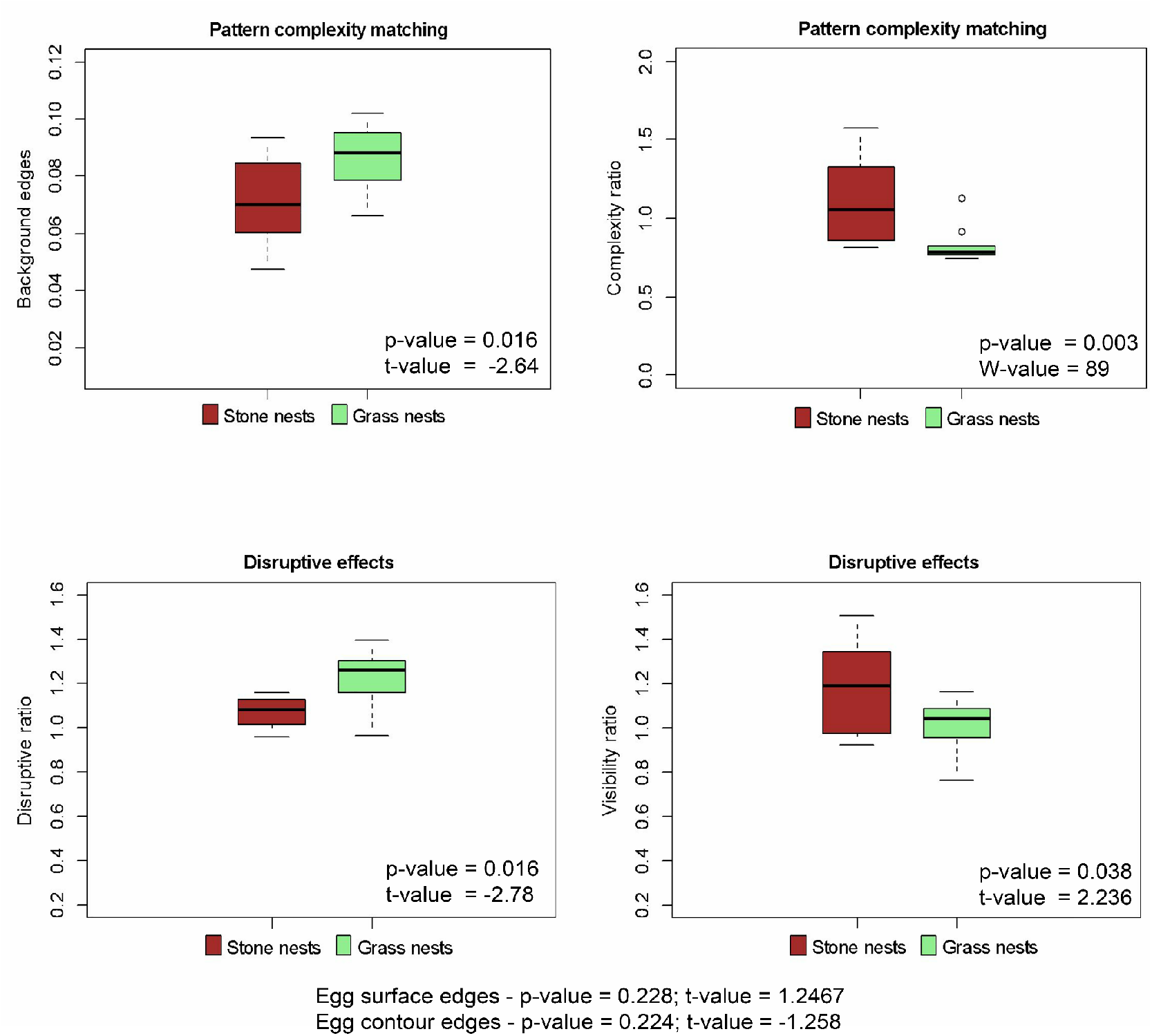
Camouflage comparison between stone nests and grass nests in terms of pattern complexity matching and disruptive effects.

The complexity ratio (CompRat), representing the ratio of interior egg edges to background substrate edges, was lower in grass nests and closer to 1 for both nest types, indicating a similar level of pattern complexity matching (Figure 8). However, it is worth noting that there was higher variability in CompRat for stone nests compared to grass nests. This suggests that grass nests may have a more consistent pattern complexity match with their surroundings, potentially contributing to their superior camouflage abilities.

#### Comparing disruptive effects

Our analysis revealed that there was no notable difference in the number of contour edges (ContEdge) between the two types of nests (p-value=0.224; t-value=-1.258). However, when we compared the disruptive ratio (DisRat), which is the proportion of edges in the egg contour area (ContEdge) to edges within the egg interior (EggEdge), we observed that the stone nest clutches were better disguised. This was because they had a significantly lower DisRat and were closer to 1 (Figure 8). These findings suggest that the edges in the egg contour region were more noticeable in grass nests than in stone nests. Whereas, the visibility ratio (VisRat), which measures the detectability of the contour edges (ContEdge) relative to the edges in the background substrate (BgEdge) was lower and closer to 1 in grass nests than in stone nests, so based on this metric, the grass nests were better camouflaged (Figure 8).

## DISCUSSION

Birds select their nest sites based on various environmental factors, such as availability of food, predator pressure, exposure to weather conditions, and interactions with other birds (Momberg et al., 2023). For Yellow-wattled Lapwings, we investigated the role of vegetation cover and food availability in their nest-site selection.

The Yellow-wattled Lapwings, like some other waders (Smart et al., 2006), preferred nest sites with higher vegetation cover that allows them to conceal their nests within the surrounding vegetation. However, this strategy may also put incubating parents at a disadvantage, as they may not be able to detect predators early through visual cues (Laidlaw et al., 2020). Despite this, vegetation cover provides shelter and protection to nesting birds from predators and environmental stresses, which may increase nesting success (Laidlaw et al., 2020). This could explain why the number of nests increased with an increase in vegetation cover in our study area. The relationship between hatching success and vegetation cover was inconclusive. Although there was a trend where the chances of successful hatching slightly increased with increasing vegetation cover, we speculate that other factors such as predator presence and anthropogenic activities could affect hatching success. Additionally, the low sample size of only 26 nests over two field seasons also makes it difficult to draw definitive conclusions about the role of habitat variables on nest site selection in this study.

We found that food availability may not be the primary factor influencing the breeding behavior of Yellow-wattled Lapwings. The presence of fewer nests in areas with high insect density points to potential competition for resources, including food and nesting sites, among the model species or other bird species (Martin et al., 1998; Newton, 1998; Krist, 2004). This observation is further supported by the higher hatching success rate of nests with lower food availability, contradicting the common belief that high food availability leads to better breeding success in birds (Li & Martin, 1991). Such sites may experience reduced competition for resources and attract fewer potential predators or disturbances from other individuals or bird species. This reduced disturbance could lower the risk of nest predation and, in turn, increase the likelihood of successful hatching. Additionally, the seasonality of food availability might play a role in these outcomes, as food availability varies across different months and habitats (Figure 9). It is essential to consider that the insects sampled may not fully represent the entire diet of the birds. The birds may be also consuming other invertebrates like millipedes (Mukherjee et al., 2015), which could contribute to the overall food abundance observed in our study. Further studies are needed to elucidate the intricate interactions between food availability, competition, and nest success in Yellow-wattled Lapwings.

**Figure 9.**
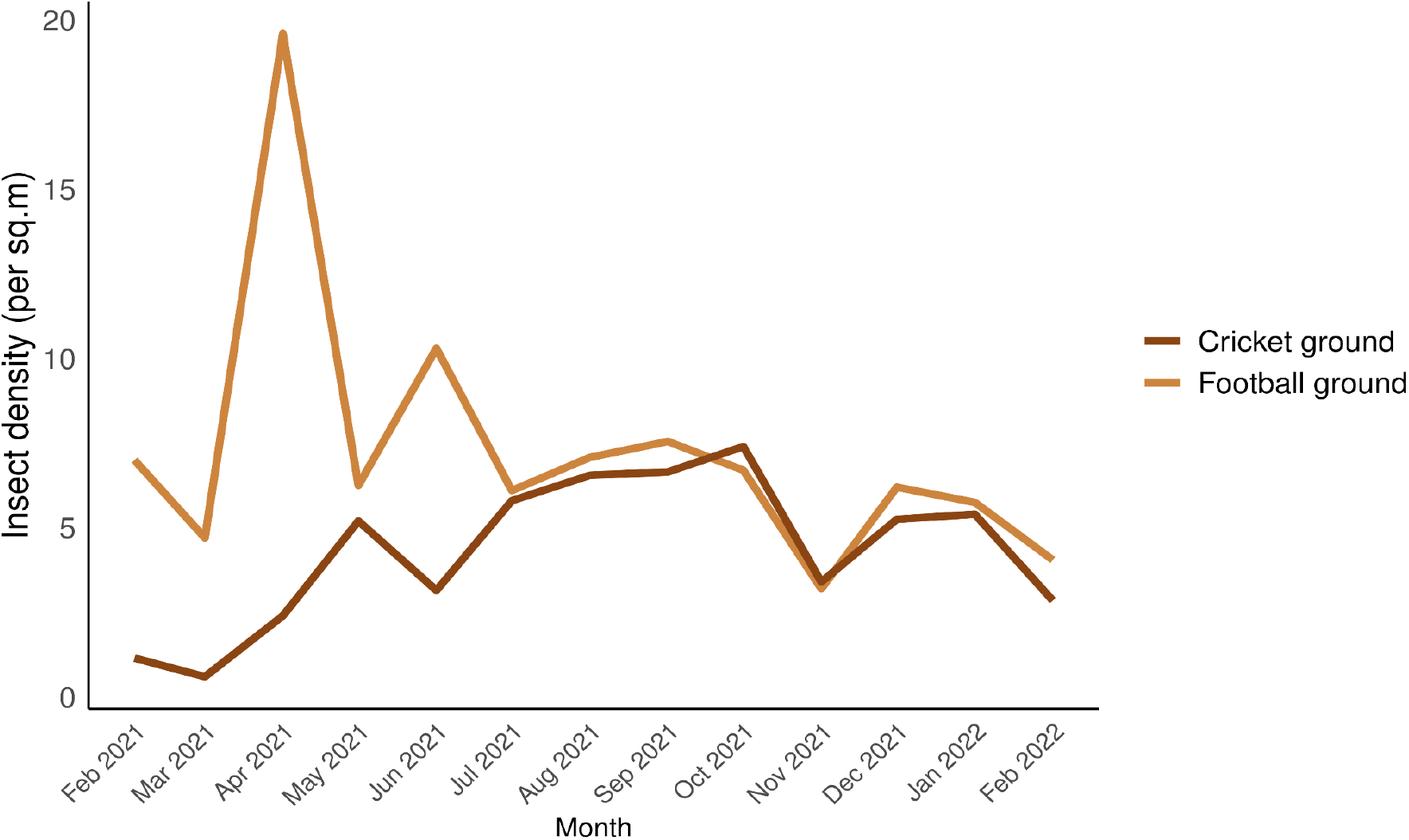
Insect density of the study sites from February 2021 to February 2022

This study is part of a growing trend in utilizing digital photography and image analysis to investigate the diverse camouflage strategies employed by animals in nature. This approach enables the comprehensive and quantitative analysis of detailed color and pattern data from both the organism and its background substrate in images, providing valuable insights into various camouflage forms (Stoddard et al., 2016).

Our analysis of different camouflage metrics and their impact on hatching success in Yellow-wattled Lapwing nests yielded intriguing results. While the complexity ratio, disruptive ratio, and visibility ratio were identified as significant factors in determining nest camouflage, statistical models did not find any camouflage metric to be a potential predictor of nest survival. This outcome raises questions about the extent to which egg camouflage explains differences in egg survival in this species, at least in the study area.

Several factors could account for this unexpected finding. Differences in visual cues evaluated by various predator species, including avian, reptilian, and mammalian animals, may contribute to the variation in egg-searching behavior. Moreover, the luminance vision and color vision mechanisms vary among predators, with non-primate mammals being dichromats (two color cones), snakes being trichromats (three color cones), and birds being tetrachromats (four color cones) (Stoddard et al., 2016). Thus, dichromatic predators may primarily rely on achromatic cues like patterns, supplemented by chromatic cues when searching for eggs (Stoddard et al., 2016). Consequently, specific interactions between achromatic and chromatic camouflage metrics could exist.

To address this complexity, quantifying additional camouflage mechanisms, such as background color matching, and conducting experimental studies to identify the significant features of camouflage for different predator species would be valuable. Such an approach would enhance our understanding of the intricate relationships between various camouflage metrics and their effects on egg survival.

Egg camouflage is an important defense mechanism against predators, but in some cases, it may not be sufficient and is likely to be the last line of defense against predators (Stoddard et al., 2011). In the case of Yellow-wattled Lapwings, visual concealment of nests through vegetation and the behavioral experience of nesting parents might have been the major contributing factors contributing to the lack of a significant relationship between camouflage and hatching success rates. The observation of aggressive behavior by parent birds towards nearby predators suggests that these experienced parents may have compensated for the suboptimal camouflage of their nests with behavioral defense mechanisms. As a result, the effects of camouflage might have been overridden, leading to no significant difference in hatching success rates between nests with different levels of camouflage. In the study (Salek & Cepáková, 2006), the researchers have found that among Charadriids, larger birds rely more on aggressive attacks as a defense strategy, while smaller birds use egg camouflage and injury feigning. They observed that Northern Lapwings primarily used aggressive attacks to defend against predators, while Little Ringed Plovers relied more on camouflage. The researchers suggested that small birds benefit more from camouflage, while larger birds use a combination of defense strategies for greater effectiveness.

In the field site, the observation of lower predation rates (7 out of 26 nests were predated) suggests that the survival of some less-camouflaged nests might have been facilitated. Predation events were categorized as direct observations or inferred based on circumstantial evidence. In a separate study by Sethi et al. (2010), they reported a predation rate of 51.73%. Given the constraints of a limited sample size, our study area exhibited an estimated predation rate (percentage of the number of nests predated out of the number of nests observed) of 26.92% (SE=0.0866). This finding implies that predation might not have exerted a significant impact on the hatching success of the eggs.

It’s important to acknowledge other influencing factors. Anthropogenic activities and parental abandonment could have influenced the study’s outcomes. Four nests were destroyed due to human activities, and one nest was abandoned by its parents. These external factors could have contributed to the overall hatching success rate of the nests, potentially overriding the effects of nest camouflage.

Our study found that different types of nests employ distinct strategies to achieve better camouflage, consistent with previous findings suggesting individual variation in camouflage mechanisms within a single species (Lovell et al., 2013). We observed that grass nests had a higher number of background edges compared to stone nests, indicating that they were laid on substrates with more edges, enhancing their camouflage (Stoddard et al., 2016). Additionally, grass nests exhibited lower complexity ratios than stone nests, suggesting better camouflage in grass nests.

Regarding disruptive effects, we found that stone nests had a lower disruptive ratio, indicating that the markings on the eggs broke up their outline and blended them into the surrounding substrate. Conversely, grass nests displayed a lower visibility ratio, indicating that the use of substrate markings effectively concealed the eggs from view, blending them into the background. Both strategies align with the notion that nests with lower disruptive ratio and visibility ratio values possess better camouflage.

Notably, there was no significant difference in egg contour region edges and egg surface edges of both types of nests. This indicates that there is only one observable egg phenotype and suggests that individuals have a behavioral preference for specific nesting substrates. The absence of observable differences in egg morphology between the two types of nests implies that small-scale habitat level adaptation to a substrate is not observable in this study site. Instead, it appears that they might be choosing their nest substrate based on behavioral preferences, such as familiarity or previous successful breeding attempts (Martin et al., 1998).

We make a further, unreported observation on Yellow-wattled Lapwings breeding activity from this study site. The breeding season for this species is expected to take place from March/April to July in India (Ali & Ripley, 1983; Sethi et al., 2010 (Haridwar district, Uttarakhand state); C. Kumar, 2015 (Ludhiana district, Punjab state)). Whereas in the study site, it has been observed from November until June/July over the past four years (2019-2022). The first recorded instance of early breeding by the Yellow-wattled Lapwing in this area (Bhubaneshwar, India) dates to 1964 when prominent scientists, Dr. J.B.S Haldane, Dr. Helen Spurway, and Dr. S. D. Jayakar reported the breeding activities of these birds from 1964 to 1968 (Jayakar & Spurway, 1965a, 1965b, 1968). According to their notes, the first nest was found in February 1964.

In seasonal environments, birds must synchronize breeding with peaks in resource availability to ensure adequate resources for the production and growth of offspring (McKinnon et al., 2012). The start of the breeding season can vary from year to year and is primarily determined by meteorological factors and food availability (Perrins, 1965; GIL-DELGADO et al., 2005). The availability of food at the study sites varied with time, but sampling across seasons is needed to understand general trends in food availability (Figure 9). Another potential explanation for the variation in breeding activities could be the demand for multi-broods within a single season.

Field observations and a note by Jayakar & Spurway (1964) suggest that Yellow-wattled Lapwings are multi-brooded species. In multi-brooding species, an earlier initiation and completion of the first brood can increase the chances of successfully completing the second brood and increase overall productivity (Vafidis et al., 2018). The early onset of breeding activities by the Yellow-wattled Lapwings in the study area is an intriguing phenomenon that requires further investigation. By understanding the underlying factors contributing to this behavior, we can gain valuable insights into the breeding ecology of this species.

In conclusion, our study has provided valuable insights into the strategies adopted by open-ground-nesting birds in selecting a suitable nesting site and protecting their nests from predators. Our research has emphasized the need to consider multiple metrics of camouflage when evaluating nest survival. We have described a complex relationship between vegetation cover, food availability, and hatching success rate in the model species. This suggests that various ecological factors affect the breeding success of these birds, and further studies are required to fully comprehend these complex relationships.

We have also identified several critical factors that should be considered when studying the survival of ground-nesting bird nests in the wild. These include parental behavior, predation rates, and anthropogenic activities. Future studies conducted in different populations and environments, with larger sample sizes and the inclusion of behavioral defense mechanisms of parents and other camouflage strategies, may provide a more comprehensive understanding of the intricate relationship between camouflage and survival in this species. In addition, by studying the behavior of birds concerning their nesting substrate preferences and nest survival, we can gain a better understanding of how they adapt to changes in their habitat and how they might respond to environmental stressors.

## ACKNOWLEDGEMENTS

We thank Veena Birudula for her assistance in the field and Dr. Mary Caswell Stoddard for providing the camouflage quantification code. We also appreciate the support from all our BioGeoSys lab members, especially Avrajjal Ghosh, Pranoy Kishore Borah, Bikash Sahoo, Ayush Parag, and Maitreya Sil.

This project was supported by intramural funding from the National Institute of Science Education and Research (NISER), Department of Atomic Energy (DAE), Government of India.

